## Supplementary material for "Hatching with Numbers: Pre-natal Light Exposure Affects Number Sense and the Mental Number Line in young domestic chicks": Table S1

**Table S1. Descriptive statistic experiment 1**. Descriptive statistics of the accuracy in selecting each of the 10 items in the Sagittal test, and in selecting the 5 items on the left (L) and the 5 items on the right (R) in the Fronto-Parallel (FP) tests of Experiment 1.

| **Experiment 1: Ordinal and Spatial cue** | | | | | | | | | |
|---|---|---|---|---|---|---|---|---|---|
| **Test** | **Hatch Condition** | **Choice** | **Mean** | **SD** | **SE** | ***n*** | ***r*** | ***P*** | **BF** |
| Sagittal | Di-chick | **1** | **17.152** | **10.579** | **2.159** | **24** | **0.651** | **0.019** | **26.604** |
| | | **2** | **15.378** | **8.408** | **1.716** | **24** | **0.682** | **0.027** | **18.426** |
| | | 3 | 11.294 | 7.015 | 1.432 | 24 | 0.227 | 1.000 | 0.497 |
| | | **4** | **28.436** | **9.374** | **1.913** | **24** | **0.878** | **< 0.001** | **>10000** |
| | | 5 | 7.986 | 5.794 | 1.183 | 24 | -0.345 | 1.000 | 0.089 |
| | | 6 | 7.326 | 6.902 | 1.409 | 24 | -0.395 | 1.000 | 0.083 |
| | | 7 | 5.914 | 5.176 | 1.057 | 24 | -0.732 | 1.000 | 0.055 |
| | | 8 | 2.546 | 3.999 | 0.816 | 24 | -0.917 | 1.000 | 0.041 |
| | | 9 | 1.678 | 2.424 | 0.495 | 24 | -0.91 | 1.000 | 0.319 |
| | | 10 | 2.292 | 3.290 | 0.672 | 24 | -0.912 | 1.000 | 0.040 |
| | Li-chick | 1 | 14.649 | 8.392 | 1.713 | 24 | 0.569 | 0.070 | 8.031 |
| | | **2** | **18.629** | **8.566** | **1.748** | **24** | **0.828** | **0.002** | **930.306** |
| | | 3 | 9.002 | 6.779 | 1.384 | 24 | -0.0943 | 1.000 | 0.135 |
| | | **4** | **36.382** | **8.164** | **1.666** | **24** | **0.88** | **< 0.001** | **>10000** |
| | | 5 | 4.974 | 4.596 | 0.938 | 24 | -0.783 | 1.000 | 0.048 |
| | | 6 | 5.914 | 6.701 | 1.368 | 24 | -0.57 | 1.000 | 0.064 |
| | | 7 | 4.583 | 6.743 | 1.376 | 24 | -0.704 | 1.000 | 0.055 |
| | | 8 | 2.511 | 4.669 | 0.953 | 24 | -0.822 | 1.000 | 0.043 |
| | | 9 | 2.094 | 2.930 | 0.598 | 24 | -0.909 | 1.000 | 0.042 |
| | | 10 | 1.261 | 2.232 | 0.456 | 24 | -0.923 | 1.000 | 4.647 |
| FP Binocular | Di-chick | 1L | 6.458 | 8.272 | 1.689 | 24 | -0.473 | 1.000 | 0.078 |
| | | 1R | 6.250 | 5.566 | 1.136 | 24 | -0.591 | 1.000 | 0.060 |
| | | 2L | 9.583 | 7.790 | 1.590 | 24 | -0.058 | 1.000 | 0.178 |
| | | 2R | 5.833 | 6.019 | 1.229 | 24 | -0.626 | 1.000 | 0.059 |
| | | 3L | 7.292 | 5.103 | 1.042 | 24 | -0.539 | 1.000 | 0.069 |
| | | 3R | 6.458 | 5.610 | 1.145 | 24 | -0.626 | 1.000 | 0.062 |
| | | **4L** | **18.958** | **10.527** | **2.149** | **24** | **0.727** | **0.005** | **168.591** |

| | | | | | | | | | |
|---|---|---|---|---|---|---|---|---|---|
| | | **4R** | **19.792** | **11.371** | **2.321** | **24** | **0.724** | **0.003** | **188.153** |
| | | 5L | 9.792 | 6.164 | 1.258 | 24 | -0.006 | 1.000 | 0.190 |
| | | 5R | 9.583 | 5.882 | 1.201 | 24 | -0.127 | 1.000 | 0.169 |
| | Li-chick | 1L | 5.095 | 5.598 | 1.143 | 24 | -0.704 | 1.000 | 0.053 |
| | | 1R | 7.350 | 6.753 | 1.378 | 24 | -0.452 | 1.000 | 0.082 |
| | | 2L | 8.023 | 7.186 | 1.467 | 24 | -0.342 | 1.000 | 0.101 |
| | | 2R | 7.164 | 6.443 | 1.315 | 24 | -0.365 | 1.000 | 0.077 |
| | | 3L | 6.576 | 5.782 | 1.180 | 24 | -0.57 | 1.000 | 0.065 |
| | | 3R | 7.785 | 7.907 | 1.614 | 24 | -0.277 | 1.000 | 0.100 |
| | | **4L** | **28.851** | **12.598** | **2.572** | **24** | **0.859** | **< 0.001** | **>10000** |
| | | 4R | 11.894 | 7.503 | 1.532 | 24 | 0.337 | 0.798 | 0.741 |
| | | 5L | 10.180 | 7.009 | 1.431 | 24 | 0.026 | 1.000 | 0.237 |
| | | 5R | 7.081 | 6.262 | 1.278 | 24 | -0.436 | 1.000 | 0.074 |
| FP Monocular Left | Di-chick | **1L** | **31.591** | **18.397** | **3.755** | **24** | **0.826** | **< 0.001** | **5702.882** |
| | | 1R | 5.716 | 9.515 | 1.942 | 24 | -0.449 | 1.000 | 0.076 |
| | | **2L** | **17.327** | **9.620** | **1.964** | **24** | **0.683** | **0.009** | **64.802** |
| | | 2R | 2.775 | 4.944 | 1.009 | 24 | -0.813 | 1.000 | 0.044 |
| | | 3L | 11.907 | 9.773 | 1.995 | 24 | 0.189 | 1.000 | 0.527 |
| | | 3R | 2.348 | 3.652 | 0.745 | 24 | -0.859 | 1.000 | 0.040 |
| | | 4L | 12.780 | 8.309 | 1.696 | 24 | 0.376 | 0.532 | 1.287 |
| | | 4R | 3.990 | 6.154 | 1.256 | 24 | -0.708 | 1.000 | 0.050 |
| | | 5L | 6.625 | 7.226 | 1.475 | 24 | -0.446 | 1.000 | 0.074 |
| | | 5R | 4.940 | 4.561 | 0.931 | 24 | -0.803 | 1.000 | 0.048 |
| | Li-chick | **1L** | **23.035** | **10.533** | **2.150** | **24** | **0.863** | **< 0.001** | **>10000** |
| | | 1R | 6.256 | 6.615 | 1.350 | 24 | -0.544 | 1.000 | 0.066 |
| | | 2L | 11.109 | 7.272 | 1.484 | 24 | 0.171 | 1.000 | 0.419 |
| | | 2R | 3.244 | 4.112 | 0.839 | 24 | -0.851 | 1.000 | 0.042 |
| | | 3L | 12.500 | 7.108 | 1.451 | 24 | 0.357 | 0.455 | 1.457 |
| | | 3R | 3.643 | 5.890 | 1.202 | 24 | -0.728 | 1.000 | 0.048 |
| | | **4L** | **22.595** | **8.736** | **1.783** | **24** | **0.876** | **< 0.001** | **>10000** |
| | | 4R | 6.297 | 6.747 | 1.377 | 24 | -0.465 | 1.000 | 0.068 |
| | | 5L | 7.578 | 7.194 | 1.468 | 24 | -0.379 | 1.000 | 0.090 |

| | | | | | | | | | |
|---|---|---|---|---|---|---|---|---|---|
| | | 5R | 3.743 | 4.773 | 0.974 | 24 | -0.803 | 1.000 | 0.045 |
| FP Monocular Right | Di-chick | 1L | 3.991 | 4.710 | 0.961 | 24 | -0.823 | 1.000 | 0.045 |
| | | **1R** | **33.744** | **16.545** | **3.377** | **24** | **0.861** | **< 0.001** | **90653.398** |
| | | 2L | 2.763 | 4.737 | 0.967 | 24 | -0.872 | 1.000 | 0.043 |
| | | 2R | 15.779 | 12.802 | 2.613 | 24 | 0.411 | 0.255 | 3.214 |
| | | 3L | 4.083 | 5.947 | 1.214 | 24 | -0.729 | 1.000 | 0.050 |
| | | 3R | 9.537 | 8.191 | 1.672 | 24 | -0.0384 | 1.000 | 0.176 |
| | | 4L | 3.746 | 3.668 | 0.749 | 24 | -0.798 | 1.000 | 0.042 |
| | | 4R | 13.417 | 8.781 | 1.792 | 24 | 0.406 | 0.398 | 1.936 |
| | | 5L | 5.033 | 5.004 | 1.022 | 24 | -0.716 | 1.000 | 0.050 |
| | | 5R | 7.908 | 6.329 | 1.292 | 24 | -0.304 | 1.000 | 0.091 |
| | Li-chick | 1L | 5.375 | 7.537 | 1.539 | 24 | -0.613 | 1.000 | 0.063 |
| | | **1R** | **23.770** | **14.388** | **2.937** | **24** | **0.765** | **0.002** | **535.661** |
| | | 2L | 4.345 | 5.663 | 1.156 | 24 | -0.768 | 1.000 | 0.050 |
| | | 2R | 13.847 | 9.447 | 1.928 | 24 | 0.419 | 0.287 | 2.233 |
| | | 3L | 4.405 | 5.204 | 1.062 | 24 | -0.759 | 1.000 | 0.048 |
| | | 3R | 11.261 | 5.861 | 1.196 | 24 | 0.293 | 0.984 | 0.591 |
| | | 4L | 5.219 | 6.356 | 1.297 | 24 | -0.594 | 1.000 | 0.057 |
| | | **4R** | **21.484** | **10.734** | **2.191** | **24** | **0.807** | **0.001** | **1842.938** |
| | | 5L | 5.994 | 4.638 | 0.947 | 24 | -0.685 | 1.000 | 0.053 |
| | | 5R | 4.300 | 4.755 | 0.971 | 24 | -0.772 | 1.000 | 0.046 |
